# NINJ1 mediates plasma membrane rupture through formation of nanodisc-like rings

**DOI:** 10.1101/2023.06.01.543231

**Authors:** Liron David, Jazlyn P Borges, L. Robert Hollingsworth, Allen Volchuk, Isabelle Jansen, Benjamin E Steinberg, Hao Wu

**Affiliations:** Department of Biological Chemistry and Molecular Pharmacology, Harvard Medical School, Boston MA, USA; Program in Cellular and Molecular Medicine, Boston Children’s Hospital, Boston MA, USA; Department of Physiology, Temerty Faculty of Medicine, University of Toronto, Toronto ON, Canada; Program in Neuroscience and Mental Health, Hospital for Sick Children, Toronto ON, Canada; Program in Cell Biology, Hospital for Sick Children, Toronto, ON, Canada; Abberior Instruments, Göttingen, Germany; Department of Anesthesia and Pain Medicine, Hospital for Sick Children, Toronto ON, Canada; Department of Anesthesiology and Pain Medicine, Temerty Faculty of Medicine, University of Toronto, Toronto, ON, Canada

**Keywords:** Ninjurin1, Ninjurin2, Inflammation, Pyroptosis, Cryo-EM, Cell rupture

## Abstract

The membrane proteins Ninjurin1 (NINJ1) and Ninjurin2 (NINJ2) are upregulated by nerve injury to increase cell adhesion and promote axonal growth in neurons. NINJ1, but not NINJ2, has also been shown to play an essential role in pyroptosis by promoting plasma membrane rupture downstream of gasdermin D (GSDMD) pore formation, as well as in lytic cell death mediated by other pathways. Recombinant NINJ1 and NINJ2 purified in detergent show irregular rings of various diameters as well as curved filaments. While NINJ1 and NINJ2 both formed ring-like structures when mixed with liposomes, strikingly, only NINJ1, but not NINJ2, ruptures liposome membranes, leading to their dissolution. Because of the better feasibility, we determined the cryo-EM structure of NINJ1 ring segments from detergent by segmenting the irregular rings into shorter fragments. Each NINJ1 subunit contains a transmembrane (TM) helical hairpin (α3 and α4) that likely mediates NINJ1 membrane localization, as well as the side-by-side interaction between adjacent subunits. There are two extracellular domain amphipathic helices (α1 and α2), among which α1 crosses over to the neighboring subunit at the outside facing surface of the ring, to link NINJ1 subunits together into chains. As such, the inner face of the rings is hydrophobic whereas the outer face of the rings is hydrophilic and should repel membranes. Live cell imaging of NINJ1-deficient THP-1 cells reconstituted with NINJ1-eGFP uncovers the pinching off of NINJ1 rings from the cell surface and the loss of NINJ1 to the culture supernatant in oligomerized forms upon inflammasome activation. Formation of rings is also confirmed by super-resolution imaging of endogenous NINJ1 using anti-NINJ1 antibody. These data suggest that membrane insertion of amphipathic helices and formation of rings with a hydrophilic outer surface underlie the mechanism for NINJ1 to pinch off membranes as if it were a nanodisc-forming amphipathic polymer, leading to membrane rupture and lysis during cell death.

## INTRODUCTION

The Ninjurin (NINJ) family comprises transmembrane (TM) proteins originally discovered as adhesion molecules important for promoting axonal growth upon injury (Araki and Milbrandt, 1996). It is now known that these proteins are widely expressed in both adult and embryonic tissues, and play important roles both within and beyond the neuronal system (Araki et al., 1997). In mammals, there are two NINJ family members, NINJ1 and NINJ2, which share conserved transmembrane regions and ∼55% overall sequence homology, but no significant homology to any other known proteins. NINJ1 and NINJ2 exhibit a predicted domain architecture comprised of an N-terminal region, an amphipathic helix and two conserved hydrophobic TM helices at the C-terminal region (Kayagaki et al., 2021). The N-terminal region was shown to mediate the cell adhesion and neurite extension function of NINJ1 (Araki et al., 1997; Kim et al., 2020). The cell adhesion function of NINJ1 is also important for progression of inflammation by promoting migration of myeloid cells to inflammatory lesions (Ifergan et al., 2011; Toma et al., 2020). NINJ1 is a target of p53, and appears to exhibit complex functions in systemic inflammation and tumorigenesis (Yang et al., 2017).

Inflammasomes are supramolecular complexes that activate caspase-1 or other inflammatory caspases, which in turn cleave and activate the pore forming protein gasdermin D (GSDMD) to induce pyroptosis, a lytic form of cell death (Chou et al., 2023; Liu et al., 2021). However, a seminal study recently showed that GSDMD pore formation is not sufficient to induce the plasma membrane rupture critical for lytic cell death and the full release of damage-associated molecular patterns (DAMP) such as lactate dehydrogenase (LDH). By contrast, NINJ1, but not NINJ2, is required for LDH release downstream of GSDMD activation in pyroptosis or other forms of lytic cell death (Kayagaki et al., 2021). This process was suggested to involve NINJ1 oligomerization on the plasma membrane in order to induce membrane rupture; however, despite the functional demonstration of its importance, the mechanism by which NINJ1 oligomerizes and elicit membrane rupture remained unclear.

During the preparation of our manuscript, Degen et al. published their structural and functional analyses of NINJ1 oligomerization and concluded that their observed NINJ1 filaments or hypothetical NINJ1 pores formed by curved filaments are the molecular entities for mediating cell membrane rupture (Degen et al., 2023). Here, our structural and cellular imaging data suggest a reverse mechanism in which NINJ1 forms rings that are hydrophobic inside and hydrophilic outside, rather than pores that have a hydrophilic conduit, to break and release themselves from membranes as lipid-filled entities analogous to nanodiscs.

## RESULTS

### NINJ1 and NINJ2 Form Rings and Filaments in Detergent

NINJ1 and NINJ2 are small proteins with molecular weights of 16.3 and 15.7 kDa, respectively (Figure 1A). To address the molecular mechanism of NINJ1 function, we overexpressed NINJ1 and NINJ2, N-terminally tagged with a polyhistidine and maltose-binding protein (His-MBP) tag cleavable by the rhinovirus 3C protease, in E. coli. We purified NINJ1 and NINJ2 by amylose affinity in the detergent Lauryl Maltose Neopentyl Glycol (LMNG), and isolated NINJ1 and NINJ2 oligomers after His-MBP removal at the heavy fractions of sucrose gradient ultracentrifugation (Figures 1B and S1A). Surprisingly, NINJ1 and NINJ2 both formed heterogenous irregular rings and curved filaments, as visualized by negative staining EM (Figure 1C). By contrast, the His-MBP tag greatly reduced the ability of NINJ1 to form oligomers (Figure S1B), likely due to a steric inhibition of oligomerization by the large tag. Similarly, we purified NINJ1 tagged at the C-terminus with an eGFP-FLAG tag expressed in the human cell line Expi293, and found that it formed similar irregular rings and curved filaments when isolated by sucrose gradient ultracentrifugation after removal of the eGFP-FLAG tag using the Tobacco Etch Virus (TEV) protease (Figures S1C and S1D).

**Figure 1.**
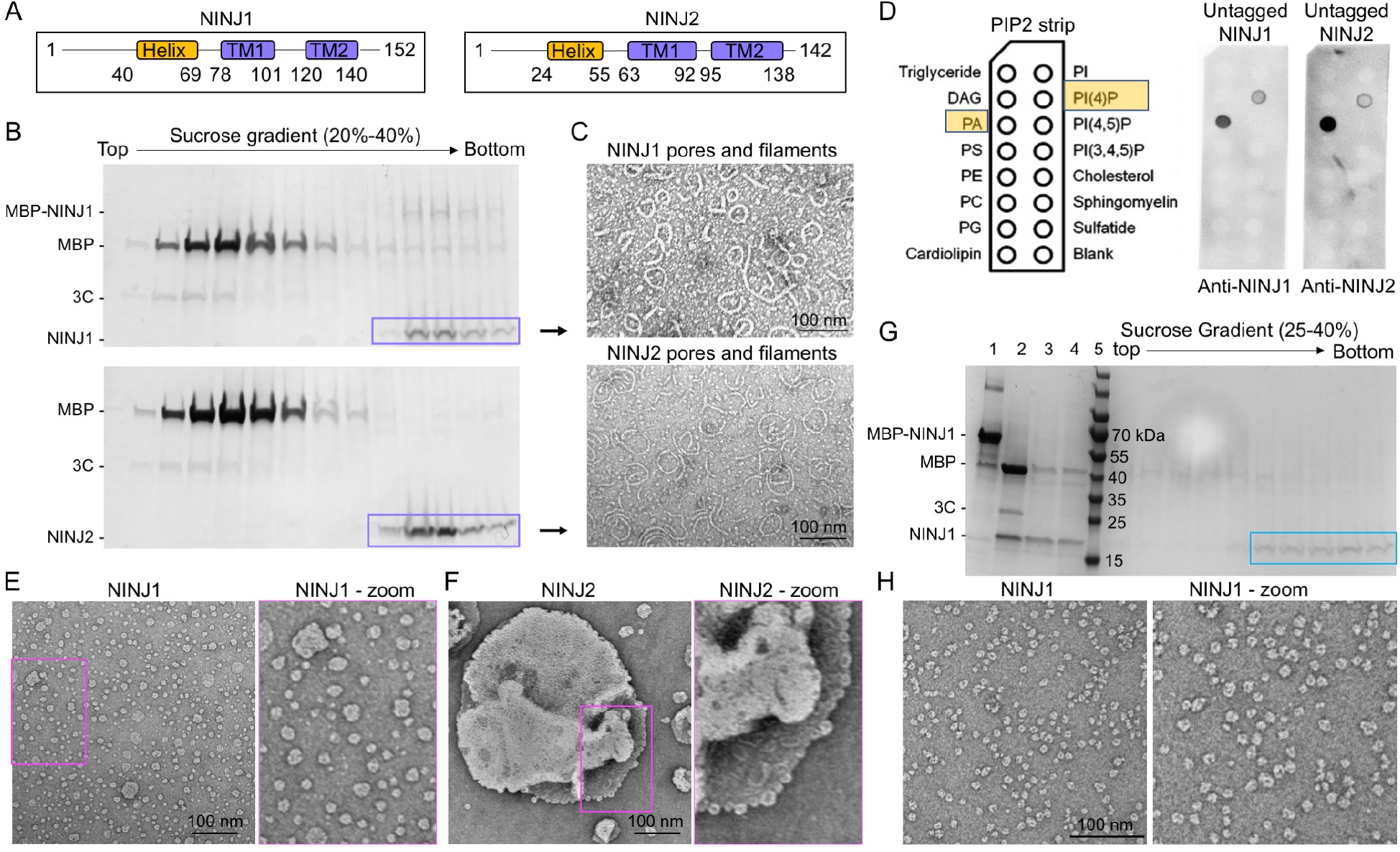
Formation of rings by NINJ1 and NINJ2, and membrane breakage by NINJ1, not NINJ2. (A) Domain architectures of human NINJ1 and NINJ2. (B) Purification of MBP-NINJ1 and MBP-NINJ2 in detergent. SDS-PAGE of NINJ1 and NINJ2 oligomers, purified from the heavy fractions of sucrose gradients after MBP removal. (C) Negative staining EM of NINJ1 and NINJ2 revealed that they both form rings in detergent. (D) Lipid strip assays for NINJ1 and NINJ2, showing that they bind negatively charged lipids, specifically PA and PI(4)P. (E,F) Incorporation of NINJ1 (E) and NINJ2 (F) into liposomes containing PA and PI(4)P, shown by negatived staining EM images and their zoom-in views. NINJ1 broke down the liposomes. (G) Further purification of NINJ1 small rings from liposomes after pelleting using sucrose gradient ultracentrifugation, shown by SDS-PAGE of the fractions (lanes at right). Lanes 1-5: His-MBP-NINJ1, cleaved His-MBP-NINJ1 by 3C, pelleted NINJ1 with lipid, pelleted NINJ1 with lipid solubilized by LMNG, and MW markers. (H) Negative staining EM of NINJ1 small rings isolated from the sucrose gradient in (G), and this same sample was used for cryo-EM studies. See also Figure S1.

### NINJ1, but not NINJ2, Breaks Down Liposome Membranes

Since NINJ1 is supposed to rupture membranes, we aimed to reconstitute NINJ1 assemblies using liposomes. For this purpose, we first identified specific lipids that NINJ1 and NINJ2 bind using lipid strip assays and recombinant, untagged NINJ1 and NINJ2 (Figure 1D). We found that both NINJ1 and NINJ2 interacted with negative charged lipids: phosphatidic acid (PA) that is the precursor of most phospholipids, and phosphatidylinositol 4-phosphate (PI(4)P), despite having contrasting isoelectric points (PI) with of 5.8 (NINJ1) and 9.5 (NINJ2). We then mixed liposomes containing phosphatidylcholine (PC, a major polar lipid), PA and PI(4)P with His-MBP-NINJ1 and the 3C protease, and incubated the mixture at 4 °C overnight to allow NINJ1 oligomerization and incorporation into liposomes upon His-MBP tag removal.

Strikingly, negative staining EM analysis revealed that most liposomes were “dissolved” by NINJ1, leaving smaller but heterogeneous structures that likely contain both lipids and NINJ1, rather than mostly large liposomes before mixing with NINJ1 (Figures 1E and S1E). Further zooming-in suggested that these small structures mimic the smaller irregular rings also observed in detergent (Figures 1C and 1E). By contrast, when we performed the same assay for NINJ2, we detected by negative staining EM irregular rings and filaments similar to the ones formed in detergent; they are however, sitting on liposomes without breaking the liposomes into small pieces as NINJ1 did (Figure 1F). Thus, these data suggest the surprising possibility that NINJ1 ruptures membranes by cutting them into small pieces, leading to lytic cell death and more complete release of DAMPs.

### Cryo-EM 2D Classes of Smaller NINJ1 Rings and Structure of NINJ1 Ring Segments

We first pursued cryo-EM structure determination of the smaller rings from liposome reconstitution. To enhance the sample quality and homogeneity, we solubilized the mixture containing liposomes and NINJ1 with LMNG followed by purification over sucrose gradient ultracentrifugation (Figure 1G), which enriched the smaller irregular NINJ1 rings formed upon mixing with liposomes (Figure 1H). 2D classification revealed rings of different sizes and shapes on top or bottom views, as well as certain details that may indicate the different subunits in the assemblies (Figure 2A); however, the heterogeneity of the data did not allow high resolution 3D reconstruction.

**Figure 2.**
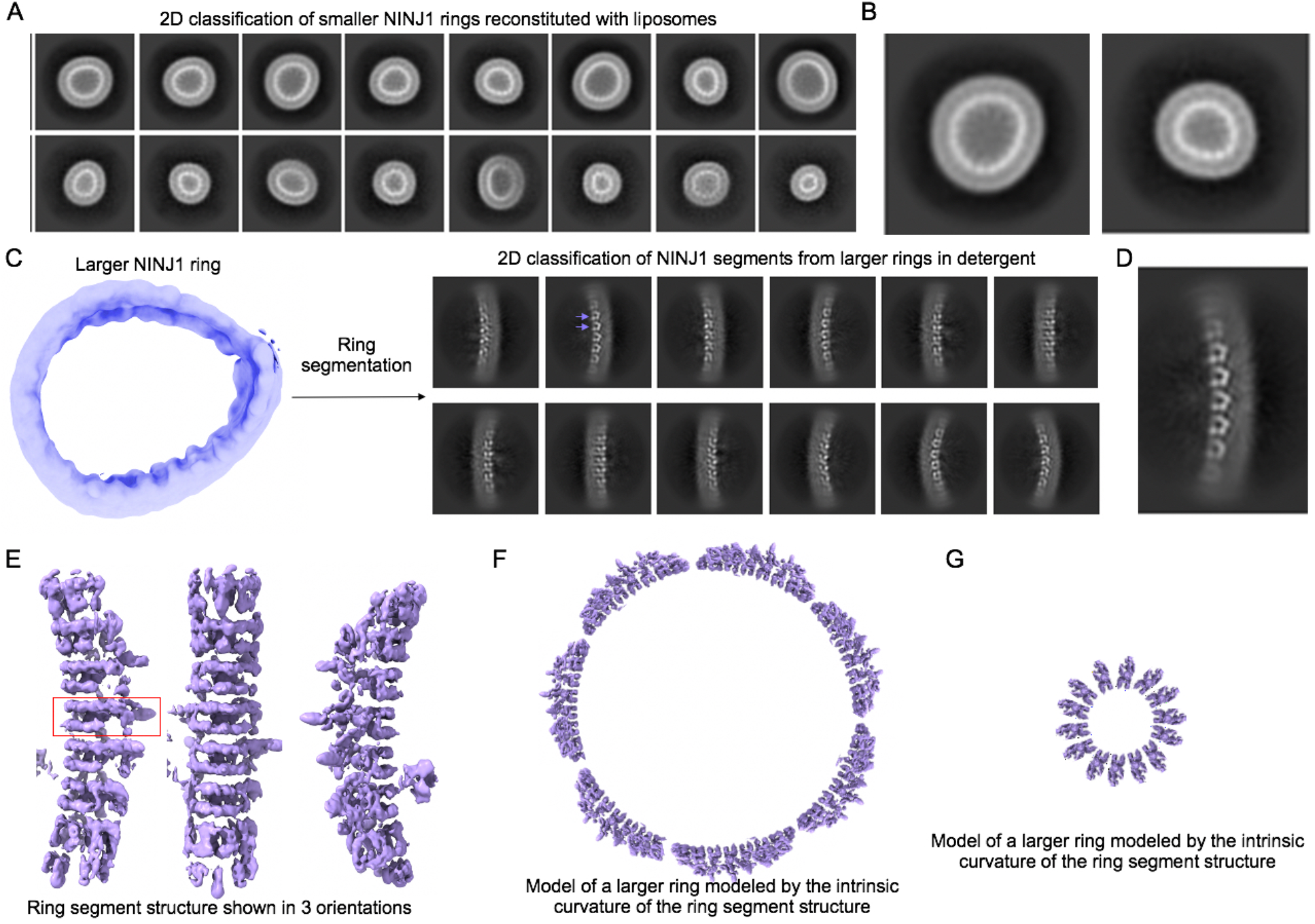
Cryo-EM studies of heterogenous NINJ1 rings and ring segments. (A) 2D classification of smaller NINJ1 rings reconstituted with liposomes, showing heterogeneous rings. (B) Magnified images of 2 classes from (A). (C) Left: irregular large rings in detergent as represented by a low-resolution 3D reconstruction. These rings were segmented into short segments as single particles. Right: 2D classification of NINJ1 segments, showing clear secondary structure details. A pair of arrows point to TM α-hairpins in two neighboring subunits. (D) A magnified 2D class of NINJ1 segment. (E) Final electron density map of NINJ1 ring segment at 4.3 Å resolution, shown in three views. (F,G) Modeled NINJ1 rings from the natural curvature of the ring segment in (E) and from an acuter curvature to construct smaller rings. See also Figure S2.

To find an alternative route to obtain a NINJ1 structure, we hypothesized that larger and smaller ring structures are assembled by similar mechanisms because they coexist in detergent, and because the smaller NINJ1 rings in detergent and from liposomes looked similar. Thus, we went on to pursue the cryo-EM structure of the larger rings in detergent. Because of the irregularity of the rings, we picked segments of these rings that later turned out to contain 7-8 subunits as single particles. 2D classification reveled potentially secondary structural details from NINJ1 TM hairpins (75-140 aa) (Figure 2B). 3D reconstruction with C1 symmetry led to a structure of the NINJ1 segment at 4.3 Å resolution (Figures 2C and S2). The natural curvature of the ring segment allowed us to generate models illustrating how rings may be formed by repeating these segments (Figure 2D), or how rings of other sizes may be formed by changing the curvature at which the neighboring subunits interacts (Figure 2E). As shown below, the inner face of these modeled rings is hydrophobic, while their outer face is hydrophilic, different from GSDMD rings (Xia et al., 2021).

Model building using the helical features of the density resulted in a subunit structure comprised of 2 amphipathic helices (37-70 aa, α1 and α2) that are roughly 90° to each other, and a pair of TM helices (78-104 for α3, 113-140 for α4) (Figures 3A). The N-terminal segment critical for cell adhesion but not pyroptosis is disordered and not seen in our structure. Our structure of the NINJ1 subunit is similar to 8CQR of Degen et al. (Degen et al., 2023), except that our TM helices are highly curved, while those in 8CQR are straight (Figure 3A). The straight helices in 8CQR might be due to the stacking between two filaments through their hydrophobic surfaces, the surfaces that face the ring center in our structure (Figure 3B). Artificial stacking may occur in vitro, especially in detergent, and our previous GSDMA3 and GSDMD structures both showed nonfunctional stacking among pores and pre-pores (Ruan et al., 2018; Xia et al., 2021). Likely due to the same reason, the filament assembly itself is also straight, rather than curved (Figure 3B), which then fails to distinguish the inner face from the outer face when it forms a ring, unlike our ring segment assembly (Figure 3C).

**Figure 3.**
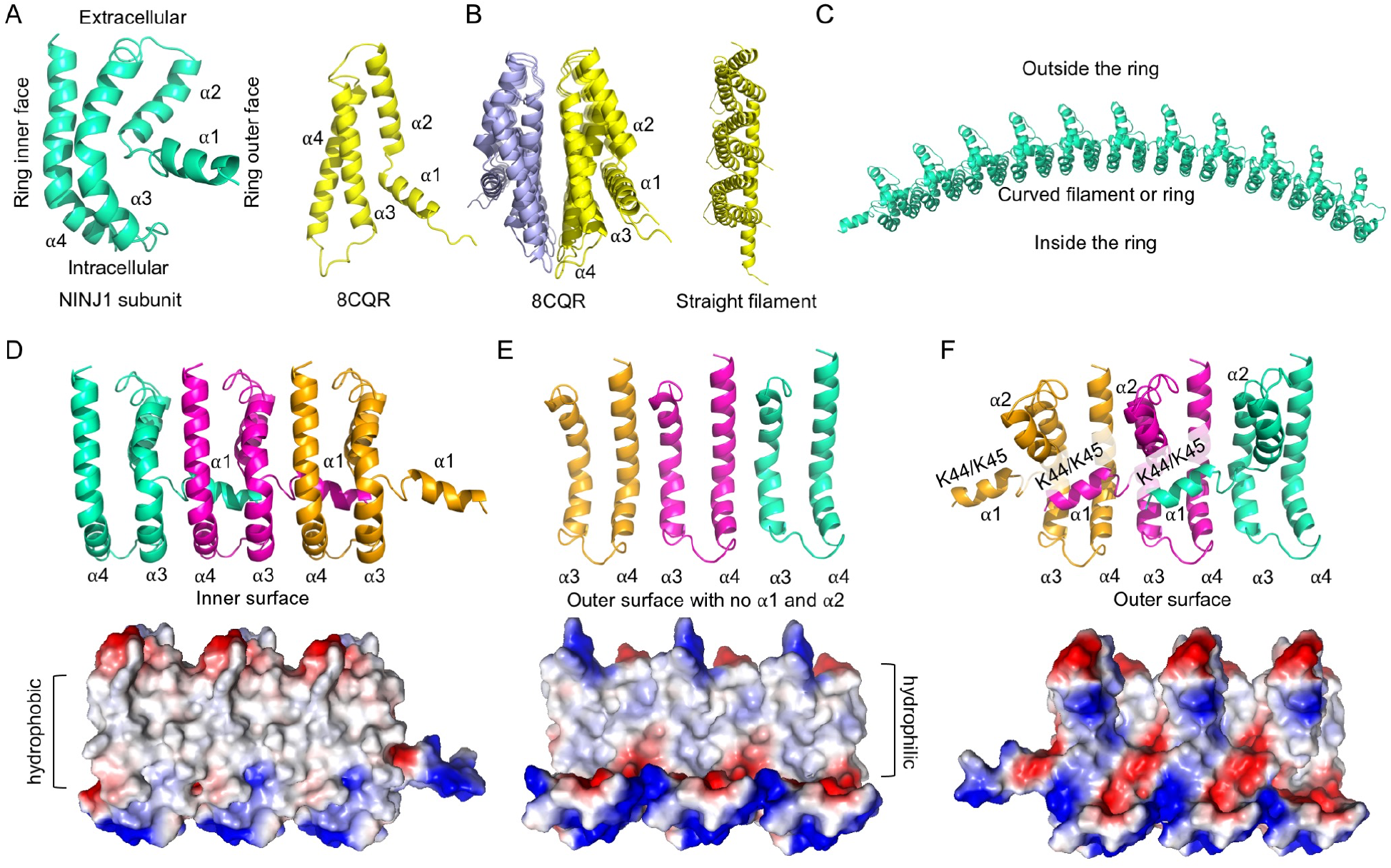
Analysis of the NINJ1 ring segment structure. (A) Cartoon representation of a NINJ1 subunit (left). α1 - α4 are labeled. Its relative orientations in a NINJ1 ring are illustrated. Cartoon representation of the published NINJ1 subunit structure (PDB ID: 8CQR) (Degen et al., 2023) is shown at right. (B) Stacking in the NINJ1 double filament structure in 8CQR, and the resulting straight TM helices and straight filament. (C) Model of a longer NINJ1 ring by propagating the ring segment oligomer, showing the clear curvature. (D-F) Ribbon representations of 3 NINJ1 subunits (top) and their electrostatic surfaces (bottom) showing the inner, lipid-facing surface of the full assembly (D), the outer, hydrophobic surface of the assembly containing only the TM helices (E) and the outer surface view of the full assembly showing the hydrophilic nature. See also Figure S3.

While the TM helices α3 and α4 pack sequentially against each other in a way that creates a curvature towards the ring center, α2 helix sits adjacent to α3 at the outer face of the ring. Importantly, α1 crosses over to the adjacent subunit, also at the outer face of the ring to promote NINJ1 oligomerization (Figures 3D). The inner surface of the ring is highly hydrophobic (Figure 3D). If we only consider the TM helices α3 and α4, the outer surface of the ring also has a hydrophobic section that could traverse the membrane (Figure 3E). By contrast, in the context of the entire structure, the outer surface of the ring is highly charged (Figure 3F). In particular, residues K44 and K45 of α1, known to be critical for NINJ1 oligomerization (Kayagaki et al., 2021), appear to contribute to inter-subunit NINJ1 interactions via their positive charges. These data support the idea that the ring is filled with membrane, whereas insertion of the amphipathic α1 and α2 both oligomerizes NINJ1 and severs the surrounding membrane, leading to release of “membrane plugs” and rupture of the membrane.

### NINJ1 Forms Rings and Sheds in THP-1 Cells during Pyroptosis

To investigate NINJ1 assemblies in macrophages for testing our structural and biochemical data, we used the human monocytic cell line THP-1 differentiated into macrophages by phorbol 12-myristate-13-acetate (PMA). We used CRISPR/Cas9 to generate clonal NINJ1 knockout cells (clones 37-39) in this background, which we validated by western blot and genomic sequencing (Figure S3). We then reconstituted these cells (clone 39) with NINJ1-GFP under a doxycycline (Dox)-inducible promoter.

Consistent with its role downstream of inflammasome activation, NINJ1 deficiency did not affect ASC speck formation upon priming with lipopolysaccharide (LPS) and subsequent NLRP3 activation with nigericin (Figure 4A). Upon Dox induction for 24 hours, THP-1 cells displayed apparent NINJ1-eGFP expression; however, NLRP3 activation by LPS and nigericin reduced the intensity of NINJ1-eGFP compared to untreated cells when imaged under the same settings (Figure 4B). The latter may indicate that some NINJ1 was liberated into the culture media. In addition to the cell membrane, NINJ1 appeared to localize in membrane extensions from THP-1 cell surfaces in LPS primed cells, as well as in NLRP3-activated cells before complete membrane rupture (Figure 4C).

**Figure 4.**
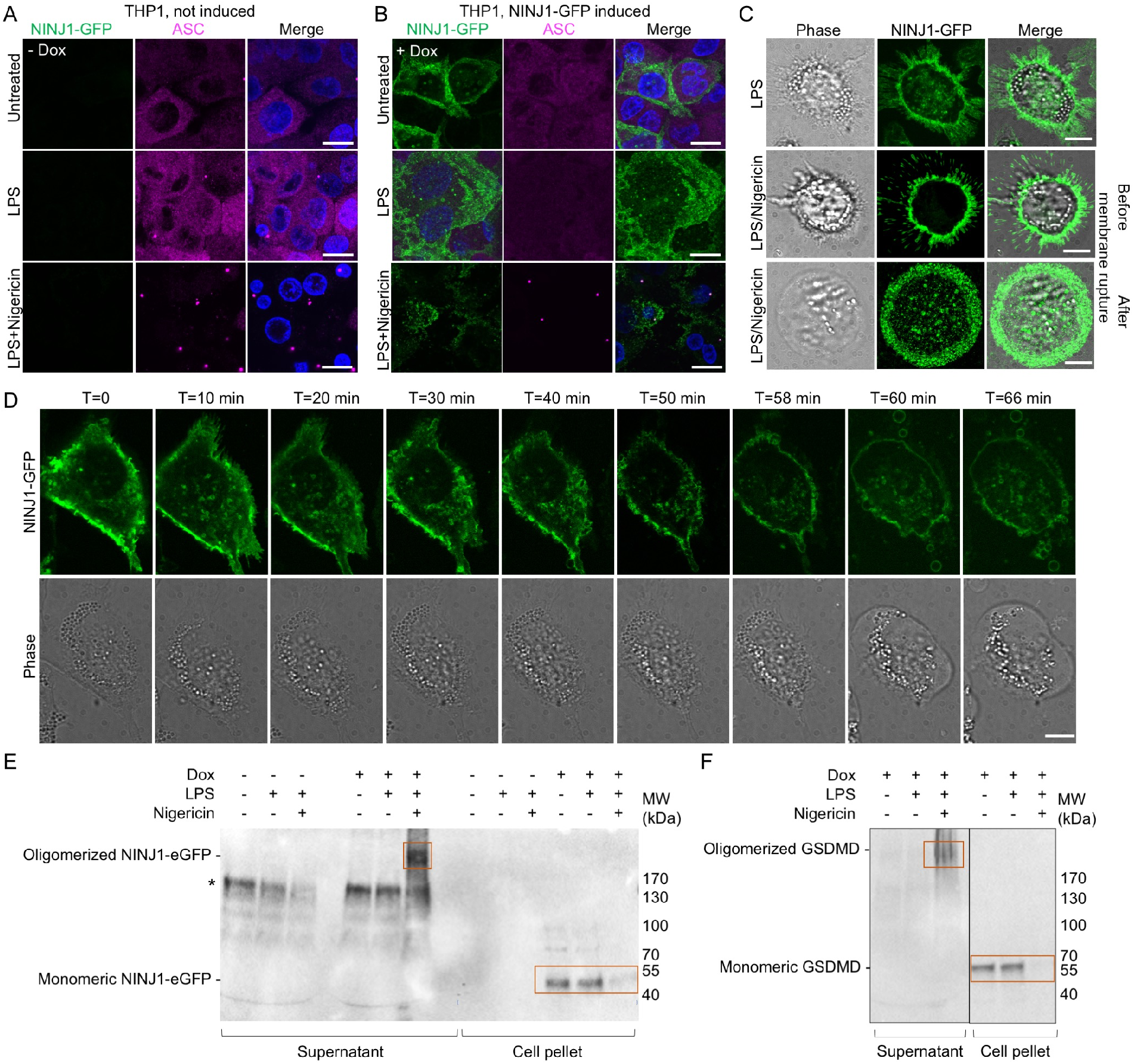
NINJ1 rings and shedding in THP-1 cells. All scale bars: 10 µM. (A,B) Fluorescence imaging of NINJ1 KO THP-1 cells reconstituted with NINJ1-eGFP for eGFP (NINJ1, green), anti-ASC (cyan), and Hoechst (DNA), for vehicle (A) or with Doxycycline induction (B). Cells were either untreated, primed with LPS (4 h) or primed with LPS (4 h) and treated with nigericin (1 h). Dox-induced cells showed NINJ1 in both untreated and LPS primed cells. After treatment with nigericin, NINJ1 expression appeared decreased. (C) Live cells fluorescence and phase images of the same THP-1 cells primed with LPS (top) or also treated with nigericin before membrane rupture (middle) and after membrane rupture (bottom). (D) Time lapse NINJ1-eGFP fluorescence (green) and phase images of the same THP-1 cells imaged for 66 min with 2 min intervals. NINJ1 shedding occurred throughout, and complete cell membrane rupture was captured between 58 and 60 min. (E,F) Western blots of NINJ1-eGFP THP-1 cell pellet and supernatant fractions using anti-NINJ1 (E) and anti-GSDMD (F). Treatment conditions for each lane are shown. Both NINJ1 oligomers and GSDMD oligomers (most likely the N-terminal fragment) were released to the supernatant after 1 h treatment with nigericin. See also Movie S1.

To further capture the live events of NINJ1 during NLRP3 activation, we recorded confocal time lapse movies of lytic cell death in Dox-induced and LPS-primed THP-1 cells upon nigericin stimulation, from time 0 to 66 min (Figure 4D, Movie S1). During this process, formation of ring-like structures of NINJ1-eGFP was observed throughout. Remarkably, we could detect the release of NINJ1-positive entities into the culture media. Some of these entities are rings with strong NINJ1-eGFP fluorescence only at the border, suggesting that they are likely NINJ1-enclosed membrane plugs, although we could not be certain that they are not some other membrane blebs generated during pyroptosis. Much smaller NINJ1 rings analogous to those observed under negative staining EM are under the diffraction limit of light microscopy and may not be seen. For the cell shown, total catastrophic membrane rupture appeared to have taken place between 58 min and 60 min, likely upon reaching a threshold in NINJ1-mediated membrane damage.

In order to determine more quantitatively the fate of NINJ1-eGFP in these cells during inflammasome activation, we performed anti-NINJ1 western blot analysis of NINJ1-eGFP in the cell pellet and supernatant fractions using non-reducing SDS-PAGE, with or without Dox induction (Figure 4E). NINJ1 was only detected in Dox-induced cells, and concentrated in the cell pellets in untreated or LPS treated cells as essentially monomers, and no NINJ1 was detected in the supernatants. Strikingly, upon nigericin stimulation, the cell pellet contained little NINJ1, and the supernatant now contained lots of oligomerized NINJ1, supporting that NINJ1-eGFP was shed to the supernatant after LPS plus nigericin treatment. We also blotted for GSDMD in Dox-induced samples. We found that while full-length GSDMD monomers occupied the cell pellets when untreated or treated with LPS alone, GSDMD was absent in the cell pellet when treated with LPS plus nigericin, and that oligomerized GSDMD (most likely the N-terminal, active fragment) was released into the supernatant (Figure 4F).

### Endogenous NINJ1 Clusters into Heterogeneous Ring-like Structures During Pyroptosis

In our previous investigations of NINJ1 regulation by the amino acid glycine (Borges et al., 2022), we conducted stimulated depletion (STED) super-resolution imaging of endogenous NINJ1 clusters in primary mouse bone marrow-derived macrophages (BMDM) stimulated to undergo pyroptosis using LPS and nigericin using anti-NINJ1 immunofluorescence. We achieved an *xy* resolution of 63 ± 8 nm by STED and demonstrated that plasma membrane NINJ1 coalesced into large clusters, following induction of pyroptosis (Borges et al., 2022). We did not observe any large clusters in LPS-primed but otherwise inactive cells, where NINJ1 was present in the plasma membrane as small punctate structures.

While we were able to visualize the noted large structures by STED, they represented the minority of NINJ1 clusters. Therefore, we next turned our attention to the major, but smaller NINJ1 clusters that could not be resolved by STED, using minimum photon fluxes (MINFLUX) nanoscopy (Schmidt et al., 2021; Tan et al., 2008). By this approach, we were able to obtain a localization precision of 2.8-4.2 nm and 3.6-7.1 nm for 2D- and 3D-MINFLUX, respectively. Representative 2D-MINFLUX images of LPS-primed BMDMs without and with pyroptosis induction are shown, as compared to standard laser scanning confocal microscopy (Figure 5A). By MINFLUX, we detected multiple NINJ1 clusters formed in cells stimulated to undergo pyroptosis, which we classified as ring-like (with a central depression of intensity), punctate (roundish, without central depression, likely limited by the diffraction limit), or other (Figure 5B). Quantification for the largest inner diameters of ring-like structures ranged from 9 to 178 nm with a median of 34.9 nm (Figure 5C), which are in keeping with the sizes of NINJ1 rings reconstituted from liposomes (Figure 1E), especially when the added dimensions from antibodies are considered. 3D-MINFLUX projection and movie for cells primed with LPS (Figure 5D, Movie S2) or treated with LPS plus nigericin (Figure 5E, Movie S3) are shown. Of note, we did not observe obvious long filament-like structures; however, we cannot exclude that some of the punctate structures are short filaments.

**Figure 5.**
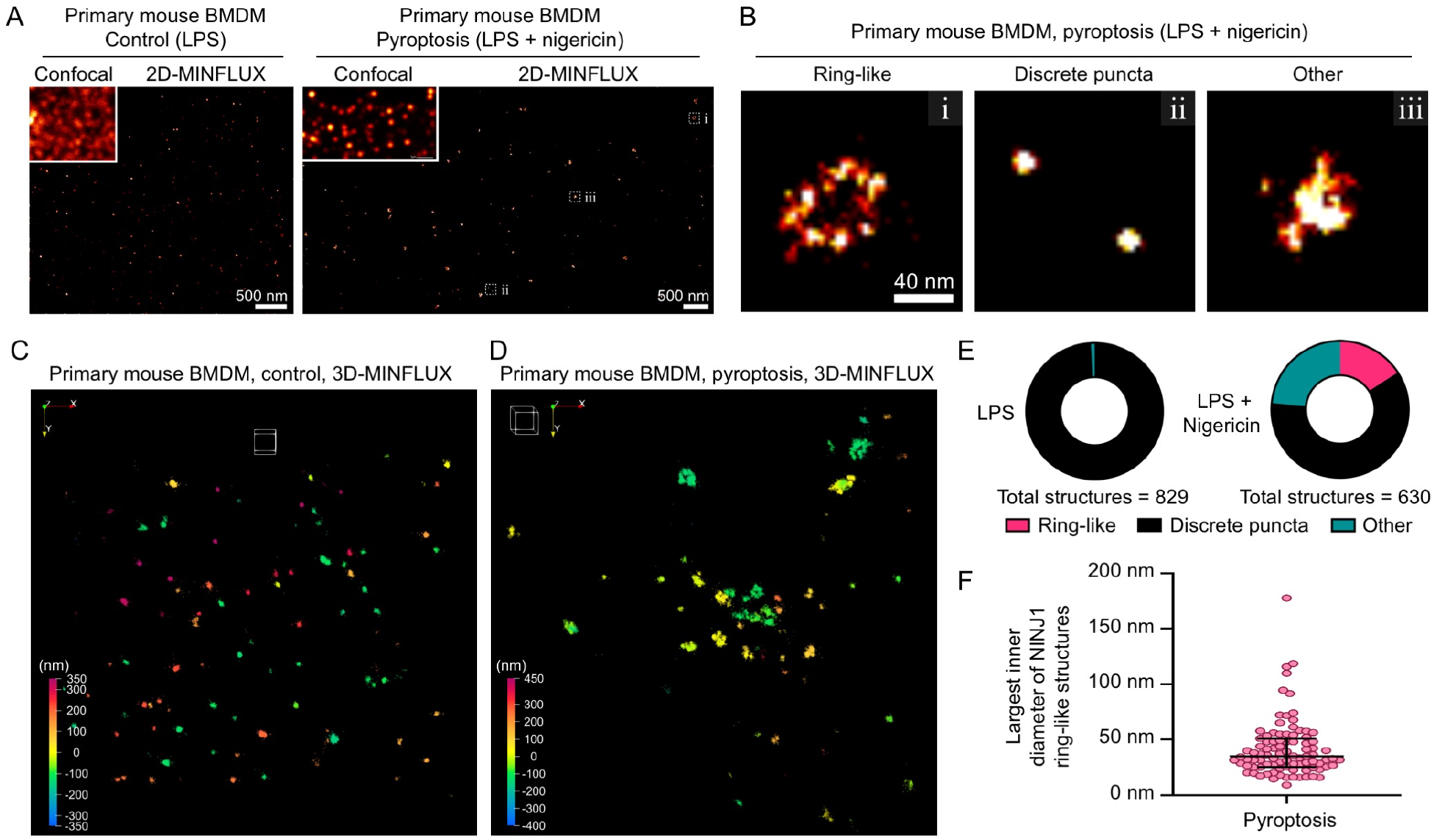
MINFLUX microscopy demonstrates that NINJ1 forms a heterogeneous population of ring-like structures during pyroptosis. LPS-primed primary mouse bone marrow-derived macrophages were stimulated to undergo pyroptosis with nigericin (20 µM, 30-60 min) or left untreated. Cells were fixed in 4% paraformaldehyde and 0.1% glutaraldehyde, followed by immunolabelling with anti-NINJ1 antibody (rabbit monoclonal clone 25, Genentech, Inc) and Alexa Fluor 647-conjugated whole IgG secondary antibody. Samples were then imaged by MINFLUX. **(A)** 2D-MINFLUX images (confocal inset) of LPS-primed macrophages without (control; left) or with pyroptosis stimulation (LPS + nigericin; right). Small dashed, white boxes indicate images shown in (B). Scale bars: 500 nm. **(B)** Representative structures of each classification category, ring-like (i), discrete puncta (ii), or other (iii). Scale bar: 40 nm. (**C-D**) 3D-MINFLUX camera-perspective renderings of heterogenous NINJ1 structures in LPS-primed cells (C) or cells stimulated to undergo pyroptosis (D). The white cube in each has an edge length of 100 nm. The rainbow-colored scale indicates distance in the Z-plane, which ranges from -350 to +350 nm (C) or -400 to +450 nm (D). **(E)** Graphs for NINJ1 structures counted and classified as ring-like (i), discrete puncta (ii), or other (iii) for LPS primed control macrophages and pyroptotic macrophages. **(F)** Largest measured inner diameter of the identified ring-like structures in the 2D plane ranged from 9 to 178 nm with a median of 34.9 nm (interquartile range 25.5, 51.0 nm). See also Movies S2 and S3.

## DISCUSSION

In this study, we used a combination of structural biology and cellular imaging to investigate the mechanism by which NINJ1 causes membrane lysis. First, we discovered that recombinant NINJ1 alone is sufficient to dissolve liposomes into heterogeneous, ring-like NINJ1-containing assemblies. By contrast, NINJ2 also formed ring-like structures when mixed with liposomes, which however failed to dissolve the liposomes. Second, we determined the cryo-EM structure of a NINJ1 assembly from segments of larger, irregular rings, whose subunit structure is similar to the published structure (Degen et al., 2023). Unlike the published structure however, which was determined from double filament bundled together by the hydrophobic side of the filament, our structure from ring segments displays highly arched individual TM helices, and an overall curvature of the assembly towards the ring center. The latter helps to determine that the inner face of the ring is hydrophobic, suitable for membrane binding. Third, live cell imaging of NINJ1 KO human THP-1 cells reconstituted with NINJ1-eGFP revealed the release of NINJ1 rings during NLRP3 inflammasome activation into the culture media, and western blot analysis showed the abundance of NINJ1 oligomers in the culture supernatant, but not in cell pellets. Fourth, super-resolution imaging of endogenous NINJ1 in mouse primary bone marrow derived macrophages identified ring-like structures of various sizes that are consistent with the irregular rings we observed in vitro. Importantly, the “chain link” connection between subunits formed by successive crossover of the α1 helix to the adjacent subunit may provide certain flexibility to accommodate different sizes and shapes, which may also depend on acidic lipid density on the membrane.

The collective data we present here illustrate the surprising mechanism on how NINJ1 may cause membrane lysis that is completely different from membrane damage and cargo release by GSDMD pores (Xia et al., 2021). This mechanism involves cutting membrane patches within NINJ1 rings, and the extraction of the membrane away from the cell surface, a process quite analogous to how certain copolymers such as styrene maleic acid (SMA) can extract membrane proteins into nanodiscs directly from cell membranes by forming a belt around the native lipid bilayer (Chen et al., 2020). We propose that NINJ1 may be upregulated by NF-κB signaling during LPS priming, as TNFR1 stimulation that also activates NF-κB has been shown to upregulate NINJ1 (Toma et al., 2020). Then, the α3 and α4 TM helices already localize NINJ1 to cell membranes in resting state, and a pyroptotic trigger then activates NINJ1, which likely allows the insertion of the α1 and α2 helices onto the outer face of NINJ1 including the crossover to generate stabilized NINJ1 rings. By acting like cookie cutters, NINJ1 rings will litter cell membranes with holes, which at a threshold point, will presumably lead to complete membrane rupture. By contrast, amphipathic filament model or pore model with a hydrophilic conduit (Degen et al., 2023) may not intuitively explain the lytic function of NINJ1, especially the release of NINJ1 oligomers into the culture media. In addition, a pore model, even if the NINJ1 pores are larger than GSDMD pores, would limit the size of its cargos, which may deter DAMP release upon lytic cell death. Thus, our data favor a cookie cutter model, if not completely exclude a filament or pore model, for NINJ1-mediated membrane lysis.

### Limitations of the study

While our study here illustrates a plausible mechanism for NINJ1-mediated membrane rupture, from which we proposed a stepwise process for this function, we do not yet know the details in these steps. In particular, our study does not address how NINJ1 gets activated upon induction of lytic cell death. One possibility is ionic influx such as Ca^2+^ upon GSDMD pore formation, but how does one explain live cell cytokine release through GSDMD pores under certain conditions (Evavold et al., 2018; Xia et al., 2021), presumably without NINJ1 activation? Another possibility stems from the unresolved characteristic large membrane bubbles in NINJ1-deficient cells upon inflammasome activation, which could suggest the involvement of membrane tension, but many alternative possibilities exist. Finally, we are intrigued by why NINJ2 oligomerizes, but does not break membranes. More studies are required to elucidate the determinant of this observation, and address other important questions related to the small but powerful NINJ1 and NINJ2 proteins.

## Acknowledgments

We thank Dr. Ed Egelman for discussions, Dr. Maria Ericsson at the HMS EM facility for training and support, Drs. Sarah Sterling, Shaun Rawson, Megan Mayer and Richard Walsh at the Harvard Cryo-EM Center for Structural Biology for cryo-EM training, help with data collection and valuable advice, Dr. Janette Myers at the Pacific Northwest Center for Cryo-EM at Oregon Health & Science University for assistance with data collection, Dr. Kangkang Song at the University of Massachusetts Cryo-EM Core for screening and preliminary dataset collection, and SBGrid team for software support and computational resources. We also thank Grigoriy Losyev at the BWH Human Immunology Center Flow Core and Ronald Mathieu at ONC-HSCI Flow Cytometry Research Facility for cell sorting and Paula Montero Llopis at the Microscopy Resources on the North Quad (MicRoN) core at Harvard Medical School for microscope use. Rabbit anti-NINJ1 monoclonal antibody (clone 25) was provided by Drs. Nobuhiko Kayagaki and Vishva M. Dixit, Genentech Inc. Graphical abstract and Figure 6 was created with Biorender.com. This work was supported by the National Institutes of Health (AI139914 to H.W.) and by a project grant to B.E.S. from the Canadian Institutes of Health Research (PJT186206).

**Figure 6.**
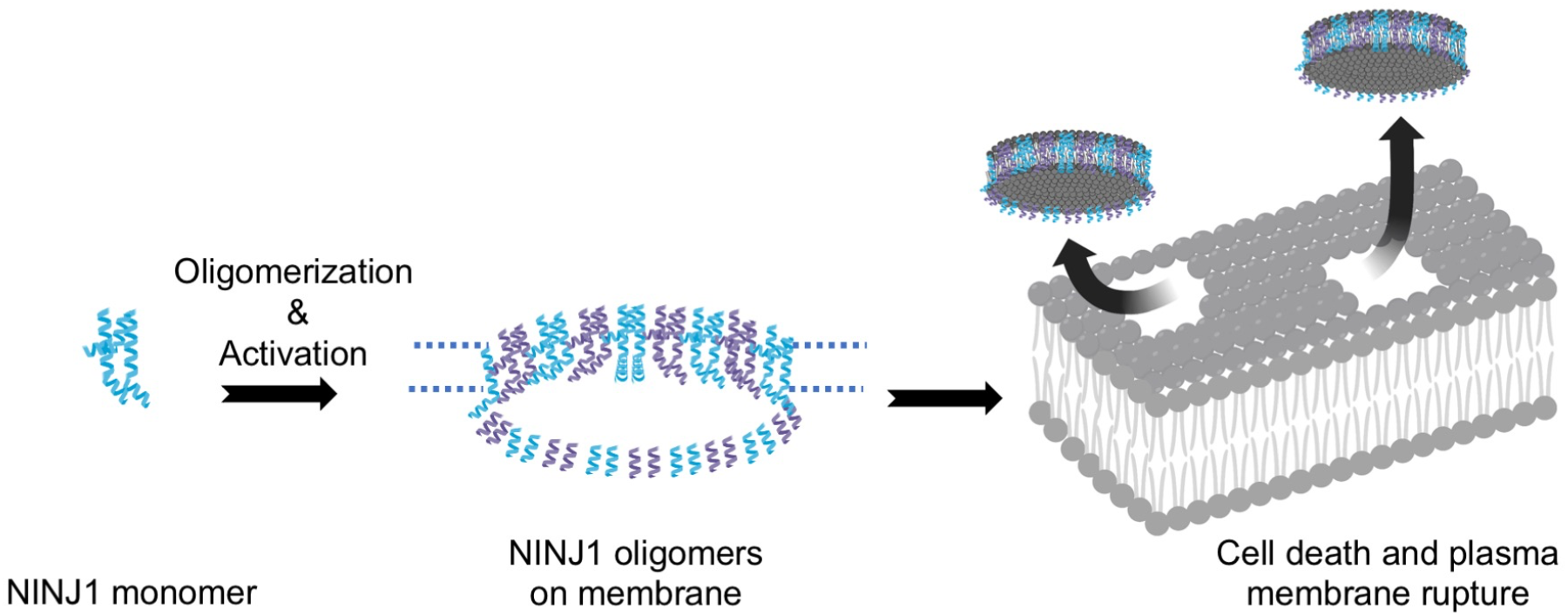
A schematic model for the proposed mechanism of NINJ1 activation and membrane rupture.

## Author Contributions

L.D. and H.W. designed and conceptualized the study. L.D. preformed cloning, protein purifications, liposome assay, negative staining EM, cryo-EM sample preparation, screening, data collection, data processing, model building and refinement. L.D. and L.R.H. generated and validated NINJ1 KO cell lines. L.R.H. cloned Dox-inducible NINJ1-eGFP construct, and L.D. generated NINJ-GFP Dox-inducible cell line. L.D. performed immunoblotting, immunofluorescence microscopy, live cell imaging, image analysis and quantifications of microscopy data. J.P.B. performed primary mouse BMDM experiments, LDH assay validation for MINFLUX sample preparation, MINFLUX data analysis and figure preparation, and contributed to manuscript writing, review and editing. J.P.B, and A.V. prepared primary mouse BMDM cultures, validation, and data curation. I.J., performed MINFLUX measurements, methodology, data curation, and MINFLUX image preparation. B.E.S. contributed to conceptualization, performed data curation, visualization, formal analysis of MINFLUX data, supervision, funding acquisition, and contributed to manuscript writing, review and editing. L.D. and H.W. wrote the manuscript with input from all authors.

## STAR★Methods

### RESOURCE AVAILABILITY

#### Lead contact

All information and requests for further resources and reagents should be directed to and will be fulfilled by the Lead Contact, Hao Wu

#### Materials availability

All requests for resources and reagents should be directed to and will be fulfilled by the Lead Contact. All reagents will be made available on request after completion of a Materials Transfer Agreement.

#### Data and code availability

- This paper does not report original code.
- Any additional information required to reanalyze the data reported in this paper is available from the lead contact upon request.

### EXPERIMENTAL MODEL AND SUBJECT DETAILS

#### Cell Lines

THP-1 (TIB-202, ATCC) cells were maintained in RPMI with GlutaMAX (Thermo Fisher Scientific, Cat. no: 11875093) supplemented with 10% FBS (Sigma), at 37 °C in 5% CO_2_ and a humidified atmosphere. THP-1 NINJ1 KO (Clones 37, 38, 39) cells were generated using the CRISPR/cas9 genome editing procedure. THP-1 NINJ1 KO cells (Clone 39) were reconstituted with codon optimized NINJ1-eGFP cloned into pLenti plasmid containing G418 selection.

### METHOD DETAILS

#### Constructs and cloning

Full-length human NINJ1 and NINJ2 were cloned into the pDB-His-3C-MBP vector with an HRV 3C protease linker for E. coli expression. In addition, full-length human NINJ1 was cloned into pcDNA3-TEV-eGFP-FLAG LIC 6D plasmid (Addgene #166835) for mammalian expression in Expi293 cells.

#### Animals and cells

Wild-type C57BL/6 animals were purchased from Jackson Laboratories (strain #000664). All animal procedures were conducted under protocols approved by the Animal Care Committee at The Hospital for Sick Children and in accordance with animal care regulation and policies of the Canadian Council on Animal Care. Mice were housed in same-sex polycarbonate cages with *ad libitum* access to food and water. Housing rooms were temperature and humidity controlled with 14:10 h light:dark cycles. Primary bone marrow derived macrophages (BMDM) were harvested from the femurs of mixed-sex cohorts of wild-type mice. Bones were then washed with PBS under sterile conditions prior to flushing the marrow by cutting the ends and centrifuging them into sterile PBS. Following a wash in phosphate buffered saline (PBS), cells were plated in DMEM with 10 ng mL^-1^ M-CSF (315-02; Peprotech Inc, Cranbury, NJ). After 5 days of culture, BMDMs were detached from the dishes with TBS and 5 mM EDTA, resuspended in fresh DMEM and plated.

#### Protein expression and purification

For E. coli expression, NINJ1 and NINJ2 constructs were transformed into BL21 (DE3) cells, grown to an OD600 of 0.6-0.8, cold shocked on ice water for 20 min, and induced overnight with 0.4 mM Isopropyl β-D-1-thiogalactopyranoside (IPTG) at 18 °C. Cells were harvested by centrifugation (4000 *g*, 20 min) and lysed by sonication in lysis buffer (25 mM Tris-HCl pH 7.5, 150 mM NaCl, 1 mM TCEP, SIGMAFAST protease inhibitor). Cell lysate was then centrifuged (40,000 *g*, 1 h); the membrane fraction was resuspended in lysis buffer supplemented with 1 % DDM: 0.1% CHS (Anatrace) and incubated for 2 h at 4 °C for extracting NINJ1 from the membrane. Following another centrifugation (40,000 *g*, 1 h), the supernatant was incubated with amylose resin for 1 h at 4 °C. Bound resin was then washed by gravity flow with 50 column volume (CV) lysis buffer with 0.1 % DDM: 0.01% CHS and eluted with 3 CV elution buffer (25 mM Tris-HCl pH 7.5, 150 mM NaCl, 1 mM TCEP, 0.1 % DDM: 0.01% CHS 50 mM maltose).

Eluted proteins from amylose resin were concentrated and cleaved overnight with 3C protease at 4 °C. These cleaved protein samples were loaded onto a step-gradient of 25%, 30%, 35%, 40% sucrose in 25 mM Tris-HCl pH 7.5, 150 mM NaCl, 1 mM TCEP, supplemented with protease inhibitor cocktail (Sigma, Cat. no: S8830) and 0.002% LMNG (x2 LMNG CMC), and ultracentrifuged for 16 h at 40,000 rpm (MLS-50 swinging-bucket rotor, Beckman). Fractions of 300 μl were collected manually from the heavy fractions of the sucrose gradient, and NINJ1-containing fractions were merged and buffer-exchanged with Zeba™ spin desalting columns (Fisher Scientific, Cat. no: PI87771) equilibrated with the buffer containing 25 mM Tris-HCl pH 7.5, 150 mM NaCl, 1 mM TCEP, supplemented with protease inhibitor cocktail (Sigma, Cat. no: S8830) and 0.002% LMNG.

For NINJ1-eGFP-FLAG expression in expi293F mammalian cells, one liter cells at 3×10^6^ cells/ml were transfected with 1 mg plasmid using polyethylenimine (3 mg/l) as transfection reagent and addition of 5 mM glycine. Cells were harvested 24 h post-transfection. They were lysed by manual homogenization in a buffer containing in 25 mM Tris-HCl pH 7.5, 150 mM NaCl, 1 mM TCEP, supplemented with protease inhibitor cocktail. The lysate was centrifuged at 42,000 RPM for 1 h (45 Ti fixed-angle rotor, Beckman), and the membrane fraction was solubilized in lysis buffer supplemented with 1 % DDM: 0.1% CHS (Anatrace) was incubated for 2 h at 4 °C, followed by another centrifugation step. The supernatant was incubated with FLAG beads over night at 4 °C. NINJ1-TEV-eGFP-FLAG was eluted with 100 μg/ml 3xFLAG peptide (ApexBio, Cat. no: A6001). Eluted fractions were concentrated and incubated with TEV protease at room temperature for 30 min, which was sufficient for eGFP-FLAG removal. The cleaved samples were loaded onto a step-gradient of 25%, 30%, 35%, 40% sucrose in 25 mM Tris-HCl pH 7.5, 150 mM NaCl, 1 mM TCEP, supplemented with protease inhibitor cocktail (Sigma, Cat. no: S8830) and 0.002% LMNG (x2 LMNG CMC), and ultracentrifuged for 16 h at 40,000 rpm. Untagged NINJ1 was isolated from the heavy fractions of the sucrose gradient.

#### *In vitro* lipid blot assay

Lipid binding assay with purified NINJ1 and NINJ2 was performed using PIP2 strip (Echelon Biosciences, Cat. no: P-6002) according to manufacturer instructions. The PIP2 strip membranes were blocked using 3% bovine serum albumin (BSA) in PNS with 0.1% Tween 20 (PBS-T) for 1 h following by incubation for 1 h with untagged NINJ1 and NINJ2 at 2 μg/ml diluted in 3% BSA in PBS-T. Next, the membranes were washed in PBS-T for 3 times (15 min total) and the bound proteins were visualized with anti-NINJ1 (R&D AF5105) and anti-NINJ2 (abcam ab172627) antibodies (1:10000 in PBS-T for 1 h). All steps were performed at room temperature. All samples were analysed at the same time under the same conditions.

#### *In vitro* liposome assay

Purified MBP-NINJ1 and MBP-NINJ2 proteins were mixed with liposomes containing 50% PC, 40% PA and 10% PI(4)P (from brain extract) at 1:3 protein: lipids ratio. The mixture was incubated for 12 h at 4 °C, together with 3C protease in order to allow MBP tag removal and NINJ1 and NINJ2 oligomerization into rings while being incorporated into liposomes. The samples were then subjected to ultracentrifugation at 40,000 rpm for 1 h for pelleting NINJ1 and NINJ2-containing fractions. The pellet fractions were visualized by negative staining EM. For cryo-EM, NINJ1 pellet fraction was further solubilized by 0.002% LMNG and subjected to sucrose gradient ultracentrifugation.

#### Negative-staining electron microscopy

For negative staining, 5 µl of NINJ1 and NINJ2 samples in detergent or from liposomes were placed on a copper grid (Electron Microscopy Sciences, cat. no: FCF400CU50), incubated for 1 min, washed twice with buffer containing 25 mM Tris-HCl, pH 7.5 and 150 mM NaCl, stained with 2% uranyl formate for 30 sec and air-dried. The images were collected at a Tecnai G2 Spirit BioTWIN or JEOL Transmission Electron Microscope (TEM) equipped with AMT 2k CCD camera at 49,000x magnification and 120 keV (HMS EM core facility).

#### Cryo-EM data collection

A 3 µl drop containing NINJ1 in LMNG (either reconstituted from liposomes as smaller rings, or detergent as mainly large rings) at 0.5 mg/ml was applied to a Lacey Carbon grid with ultrathin carbon support (Ted Pella, Cat. no: 01824G), blotted for 4 s, plunged into liquid ethane, and flash frozen using a FEI Vitrobot Mark IV at 100% humidity and 4 °C. Grid conditions were optimized during screening and small data set collection at the Pacific Northwest Center for Cryo-EM at Oregon Health & Science University (PNCC), the University of Massachusetts Cryo-EM Core (UMASS) and the Harvard Cryo-EM Center for Structural Biology (HMS) using FEI Talos Arctica (ThermoFisher) microscopes equipped with an autoloader (200 keV, Gatan K3 direct electron detector).

Final datasets were collected at HMS using a Titan Krios microscope (ThermoFisher) operating at 300 keV and equipped with a BioQuantum Imaging Filter (Gatan) and K3 direct electron detector (Gatan). Automated data collection was performed using SerialEM software (Mastronarde, 2005), and the movies were obtained in counting mode at 105,000x magnification (0.825 Å/pix). For NINJ1 smaller rings purified from liposomes we collected 11,000 movies with 48 frames each, recorded at multiple defocus values from -1.2 to -2.4 μm and with multiple exposures per stage shift (5×4) introduced with image shift. Each movie was acquired at a dose rate 27.44 e/s per physical pixel and a total accumulated dose of 51.3 e/Å^2^ over 1.31 s total exposure time. For NINJ1 sample with large rings purified in detergent, 15,296 movies with 47 frames each were recorded at multiple defocus values from -1 to -2.2 μm and with multiple exposures per stage shift (5×4) introduced with image shift. Each movie was acquired at a dose rate 12.588 e/s per physical pixel and a total accumulated dose of 52.08 e/Å^2^ over 2.8 s total exposure time. Both data sets have 0.825 Å pixel size.

#### Cryo-EM data processing

Cryo-EM data processing software and support was provided by SBGrid consortium (Morin et al., 2013). Raw movies were corrected by gain reference and beam-induced motion and combined into a motion-corrected micrograph using the MotionCorr2 algorithm (Zheng et al., 2017). The defocus value for each micrograph was determined with CTFFIND4 (Rohou and Grigorieff, 2015). Automated particle picking was performed in CryoSPARC (Punjani et al., 2017): helical picker for large ring segments, and Topaz training and automated picking for NINJ1 small ring data set.

The dataset of NINJ1 smaller rings yielded 597,920 particles, which were extracted with no binning resulting in 0.825 Å pixel size. Several rounds of 2D classifications were applied, showing preferred top and bottom views of NINJ1 in rings of various sizes and shapes.

A dataset of NINJ1 larger rings yielded 12,907,558 ring segments. 3 rounds of 2D classification resulted in a particle stack of 626,231 segments. 1 initial model was generated with *ab initio* reconstruction. Helical refinement mode, followed by the non-uniform 3D refinement using C1 symmetry resulted in final cryo-EM maps at 4.3 Å resolution. The described cryo-EM workflow for this dataset is presented in Figure S2 with gold-standard Fourier shell correlation (FSC). Post-processing of the map was performed with DeepEMhancer (Sanchez-Garcia et al., 2021).

#### Model building and structure representation

The atomic model was built using cryo-EM map obtained from the NINJ1 ring segments purified in the presence LMNG (4.3 Å resolution). Model building was performed using UCSF-Chimera (Pettersen et al., 2004) and Coot (Emsley and Cowtan, 2004). The final model was further subjected to refinement in Phenix (Adams et al., 2010; Klaholz, 2019) with the starting model as a reference. Structure representations were generated using ChimeraX (Goddard et al., 2018) and Pymol (Delano, 2002).

#### Generation of NINJ1 KO and NINJ1-eGFP cell lines

To generate NINJ1 KO and sgRNA control THP-1 cells, we transduced THP-1s (early passage, ATCC) with lentiCRISPR v2 (NINJ1 Genescript guide #6 GGCACATAGAAGGCGAAGCT, addgene #125836). For lentivirus preparation, 0.5×10^6^ HEK293T cells in 6-well dishes were transfected (6 µL Fugene) with 1 µg of plasmid containing the construct of interest, 750 ng psPAX2 packaging plasmid and 250 ng pMD2.G envelope plasmid (Addgene plasmids #12260 and #12259). On the following day the medium was changed, and the virus-containing medium was collected an additional 24 h later. Virus-containing medium was concentrated with LentiX concentrator (Takara) and resuspended in RPMI (20X concentration). 1 million THP-1 cells seeded in 6-well plates were spinfected with 100 µL concentrated virus in 900 µL medium (2 h, 30 degrees, 1000 x g) containing 8 ug/mL polybrene. Following spinfection, THP-1 cells were recovered in fresh medium for 2 d prior to puromycin selection (3 µg/mL) for 3 days. Selected cells were single cell sorted into 96 well format with a BD FACSAria II cell sorter equipped with 100 µm nozzle. Cells were expanded before screening clones by western blot and subsequent genomic sequencing. We obtained 3 clones (37,38,39) in which NINJ1 was undetectable by western blot (Figure S4). Among the 3 clones, we used clone 39 for subsequent experiments given it had biallelic frameshift mutations (1 and 11 bp, Figure S4). For reconstituting NINJ1 KO cells with doxycyling-inducible NINJ1-eGFP (pInducer20), we produced virus and spinfected NINJ1^-/-^ THP-1 cells as above. Cells were selected with 250 μg/ml G418 for 2 weeks prior to use in further experiments.

#### Immunoblotting of whole cell lysates

NINJ1-eGFP cells were seeded at 0.7×10^6^ cells/well on a 6-well tissue culture plate and treated with 25 ng/ml phorbol 12-myristate 13-acetate (PMA, P1585, Sigma) for 2 days and then recovered with complete medium (without PMA) for 24 h and supplemented with 1 μg/ml Doxycycline (Dox) or no Dox as a control. On the next day, cells were untreated or primed with 1 μg/ml LPS (Invivogen, Cat. no: tlrl-b5lps) for 4 h, followed by 1 h incubation with 10 μM nigericin (Sigma-Aldrich, Cat. no: N7143-5MG) for cell death stimulation. While collecting cells, we collected both pellet and supernatant fractions. The pellet fractions were collected by scraping cells with cell lifter with PBS and spinning at x 500 g for 5 min. The supernatant fractions were first spun down at 500 g for 5 min and then concentrated using 10K cutoff Centricon. SDS sample buffer was added to equal number of cells (by volume) and loaded to 4-20% non-reducing gel for western blot analysis. Western blotting was performed using anti-NINJ1 (1:1000, R&D AF5105) and anti-GSDMD (1:1000, Novus Biologicals, NBP2-33422). Anti-sheep-HRP (1:5000, Cell Signaling, Cat. no: 7076S) and anti-rabbit-HRP (1:5,000, Cell signaling, Cat. no: 7074S) secondary antibodies were used.

#### Immunofluorescence (IF)

NINJ1-eGFP THP-1 cells were plated on CELLview 4-compartment dishes (Greiner Bio-One), treated with 25 ng/ml phorbol 12-myristate 13-acetate (PMA, P1585, Sigma) for 2 days, and then incubated for 24 h with complete medium without PMA and supplemented with 1 μg/ml Doxycycline (DOX) or without DOX as a control. Cells were untreated or primed with 1 μg/ml LPS (Invivogen, Cat. no: tlrl-b5lps) for 4 h, followed by 1 h incubation with 10 μM nigericin (Sigma-Aldrich, Cat. no: N7143-5MG). Cells were fixed in 3% paraformaldehyde (PFA) for 30 min at 4 °C and permeabilized with 0.1% Triton X-100 for 10 min. Cells were incubated in PBS-Tween containing 3% BSA for 3 h, which minimized non-specific binding. After three washes with PBS-Tween, cells were incubated overnight at 4 °C with primary antibodies (human ASC 1:1000, Novus Biologicals, NBP1-78977). NINJ1 was detected by the eGFP signal and no antibody was required in order to enhance the signal. After incubation, cells were washed and incubated with AlexaFluor647-conjugated anti-rabbit IgG (1:750, ThermoFisher, Cat. no: A27040) for 1 h at room temperature, washed with PBS (3 × 10 min) and then stained with Hoechst (1:500, Immunochemistry Technologies, Cat. no: 639). Cells were imaged using a Nikon Ti inverted microscope fitted with a Photometrics CoolSNAP HQ2 Peltier cooled CCD camera and Andor Zyla 4.2 sCMOS camera at x60 magnification using Plan Apo 60x/1.3 DIC objectives. Lumencor SpectraX LED illuminator was used. Chroma 49000 (DAPI), Chroma 29002 (green) and Chroma 49011 (far red) filter cubes were used. Image analysis was performed in Fiji (Schindelin et al., 2012).

#### Live cell imaging

NINJ1-eGFP THP-1 cells were cultured in RPMI supplemented with GlutaMAX (Thermo Fisher Scientific, Cat. no: 11875093) supplemented with 10% FBS (Sigma) and neomycin (G418), at 37°C in 5% CO2 atmosphere. Cells treatment was carried out as for IF (above). For performing live cell imaging, we imaged LPS primed cells, induced with DOX for 24 h. Cells were imaged using a Nikon Ti inverted microscope fitted with a Photometrics CoolSNAP HQ2 Peltier cooled CCD camera and Andor Zyla 4.2 sCMOS camera at x60 magnification using Plan Apo 60x/1.3 DIC objective. Green and bright field channels were used. Time lapsed imaging was started upon addition of 10 μM nigericin. Images were recorded for over an hour with 2 min intervals. Image analysis was performed in Fiji (Schindelin et al., 2012).

#### Pyroptosis induction

Pyroptosis was activated in primary BMDMs primed with (0.5 µg/ml) LPS from *E. coli* serotype 055:B5, which was first reconstituted at a stock concentration of 1 mg/ml. Cells were primed with LPS for a total of 5 h, during which pyroptosis was induced with 20 µM nigericin (Sigma N7143; stock 10 mM in ethanol) for the final 30 or 60 min.

#### LDH assay

Cells were seeded at 300 000 cells per well in 12-well plates, treated as indicated, and cytotoxicity assayed by LDH release assay the following day. Following treatment, the cell culture supernatants were collected, cleared of debris by centrifugation for 5 min at 500 x *g*. The cells were washed once with PBS then lysed in lysis buffer provided in the LDH assay kit (Invitrogen C20300). Supernatants and lysates were assayed for LDH using an LDH colorimetric assay kit as per the manufacturer’s instructions (Invitrogen C20300).

#### Minimal photon fluxes (MINFLUX) nanoscopy

Primary cells from wild-type animals were harvested as described above and cultured on 1.5H glass coverslips. Cells were primed with LPS and stimulated to undergo pyroptosis as described above. Pyroptosis induction was confirmed by LDH release for each experimental replicate. Following treatments, cells were washed with PBS and fixed in 4% paraformaldehyde and 0.1% glutaraldehyde in 0.1 M cacodylate buffer at room temperature for 15 min. The cells were washed 3 times in PBS, permeabilized with 0.1% Tween-20, then blocked in 10% donkey serum and 0.1% Tween-20 in PBS for 1 h in PBS. Rabbit monoclonal anti-mouse NINJ1 primary antibody (clone 25; kind gift from Dr. Kayagaki and Dr. Dixit at Genentech, Inc.) (Kayagaki et al., 2021) at 10 µg/mL was then added overnight at 4 °C. The cells were washed three times with PBS supplemented with 0.1% Tween-20 before the addition of secondary antibody in PBS with 1% donkey serum for 1 h at room temperature. Alexa Fluor-647-conjugated donkey anti-rabbit secondary antibody (711-605-152) was used at a 1:1000 dilution. The samples were then washed with PBS and stabilized for MINFLUX imaging using gold nanorod dispersion (A12-40-980-CTAB-DIH-1-25, Nanopartz, Inc.) for 5-10 min, then mounted in imaging buffer as previously described (Balzarotti et al., 2017; Grabner et al., 2022).

A MINFLUX microscope (Abberior Instruments) equipped with a spatial light modulator-based beam shaping module and an electro-optical detector-based MINFLUX was used. Microscopy and MINFLUX measurements were performed using Imspector Software (v.16.3.15620-m2205-win64-MINFLUX) using MINFLUX sequence templates seqIIF and DefaultIIF3D for 2D- and 3D-MINFLUX measurements as previously described (Ostersehlt et al., 2022). Before starting each MINFLUX measurement, the stabilization of the microscope was activated. Measurements were conducted with a stabilization precision of < 1 nm. A 642 nm (CW) excitation laser, a 405 nm activation laser, and detection windows in the range of 650-750 nm were used. The emitted fluorescence photons were counted using two avalanche photodiodes with the appropriate fluorescence filters. Molecular precision was enabled through a reflection-based stabilization unit based on a 980-nm laser as previously described (Schmidt et al., 2021). For structure counting and ring-like structure diameter measurements, localizations were exported as 2D projections with a pixel size of 4 nm (2D) or 5-6 nm (3D) using Imspector Software. Individual structures were counted in ImageJ and classified according to shape: ring-like, punctate, or other. The diameter of each ring-like structure was considered as the largest measurable distance between points with zero localizations in the structure center and measured using the line tool in ImageJ-Fiji Software.

