## Supplemental data for "NINJ1 mediates plasma membrane rupture through formation of nanodisc-like rings"

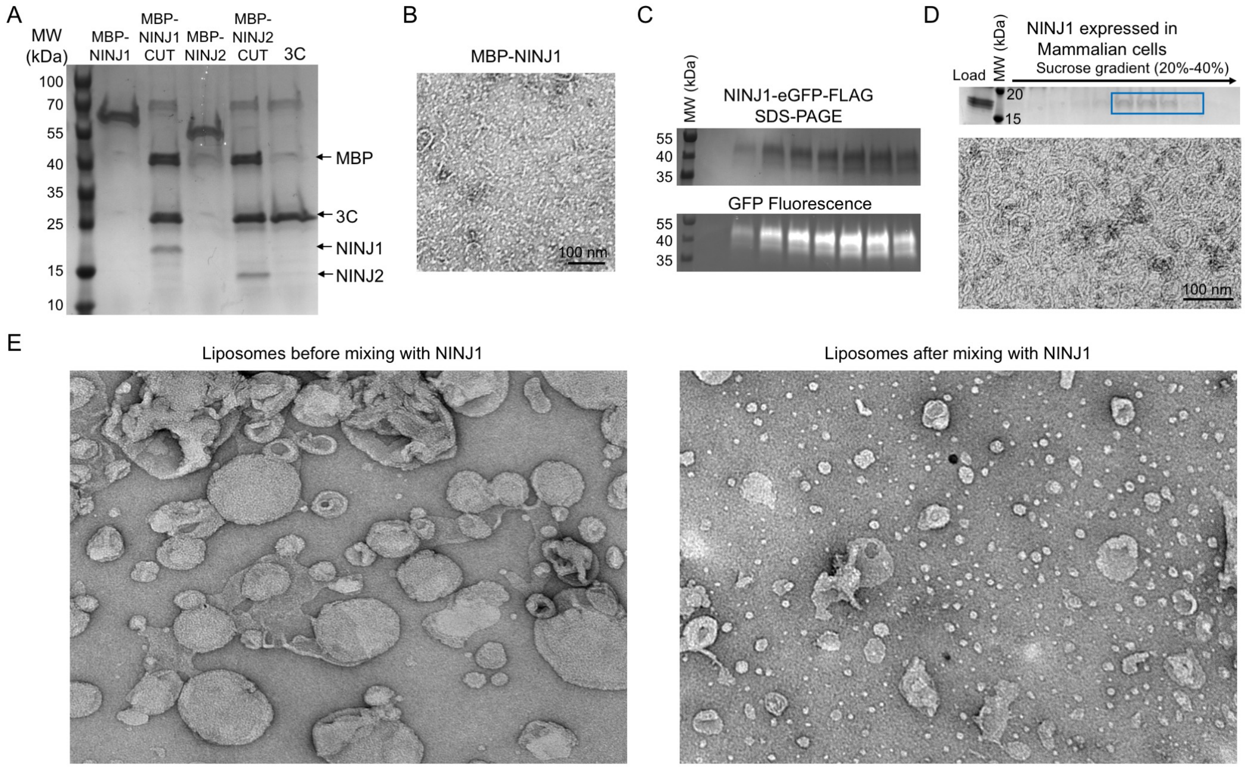


### Figure S1. NINJ1 and NINJ2 purification

(A) SDS-PAGE of MBP-NINJ1 and MBP-NINJ2 purification from E. coli expressed.
(B) Negative staining EM image of MBP-NINJ1. NINJ1 large rings were hardly detected before MBP removal.
(C) SDS-PAGE of NINJ1-eGFP-FLAG purification and detection by in-gel GFP fluorescence at the expected molecular weight.
(D) SDS-PAGE (top) and negative staining EM image (bottom) of NINJ1-eGFP-FLAG after eGFP-FLAG removal by TEV.
(E) Liposome breakdown by NINJ1, shown by negative staining EM images before and after NINJ1 incorporation into liposomes. Liposome membrane breakdown by NINJ1 is clearly shown.
Related to Figure 1.

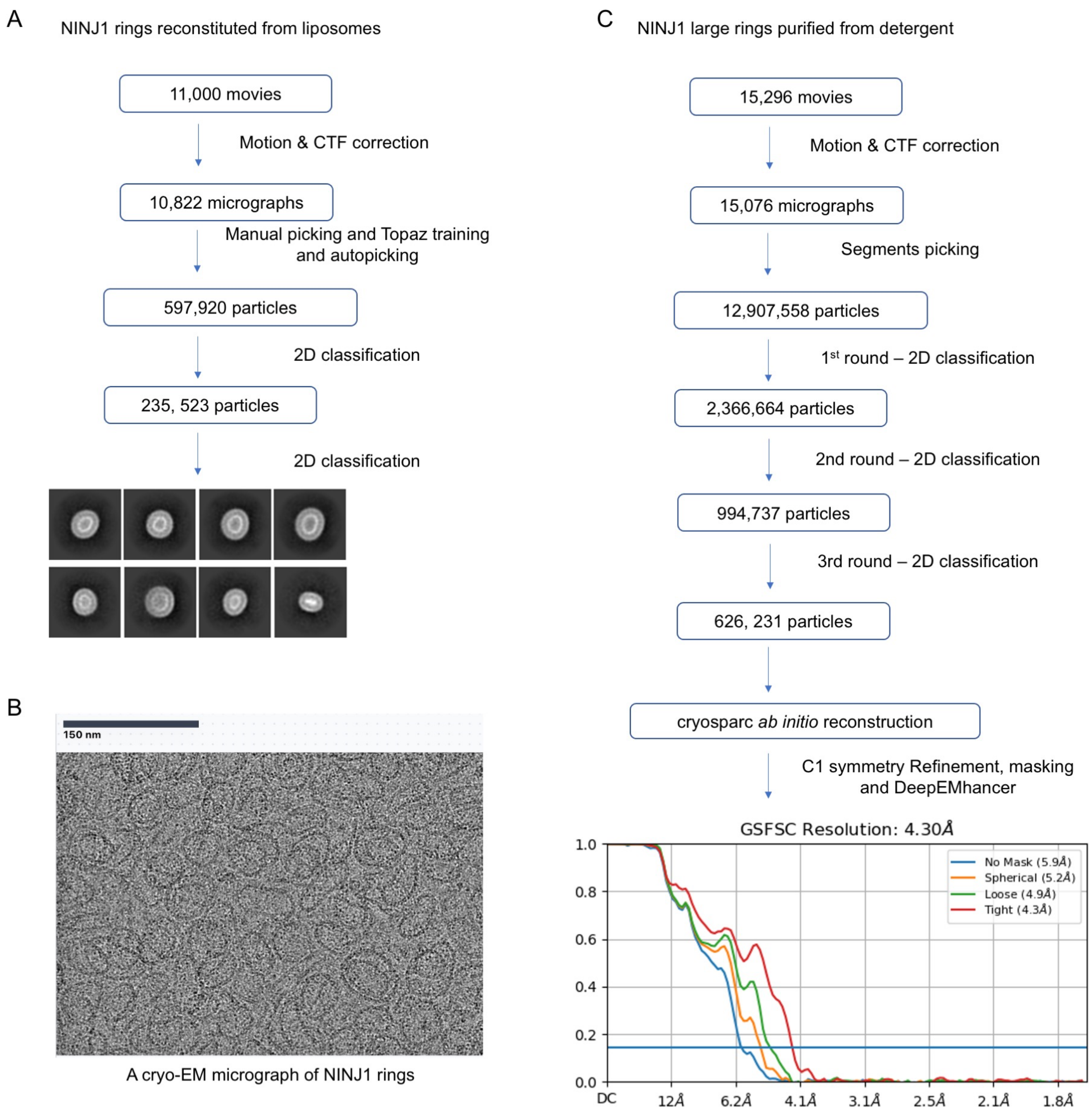


**Figure S2. Cryo-EM flow charts of NINJ1 data processing**

(A) Flow chart for cryo-EM data processing of NINJ1 rings reconstituted from liposomes.

(B) A raw cryo-EM micrograph of NINJ1 large rings in detergent.

(C) Chart-flow for cryo-EM data processing of NINJ1 large ring segments purified from detergent.

Related to Figure 2.

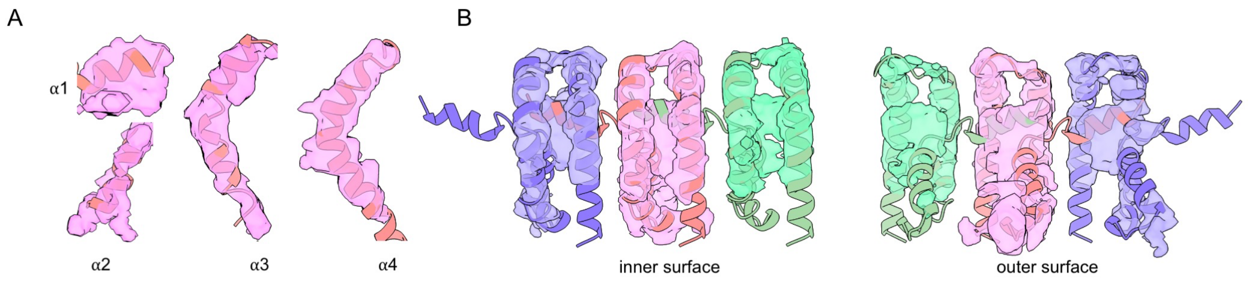


**Figure S3 – Structural representation on NINJ1 ring segments**

Different views of NINJ1 subunits fitted in the cryo-EM map.

(A) Individual helices.

(B) Side of the segment facing the inner side of the ring.

(C) Side of the segment facing the outer side of the ring.

Related to Figure 3.

A

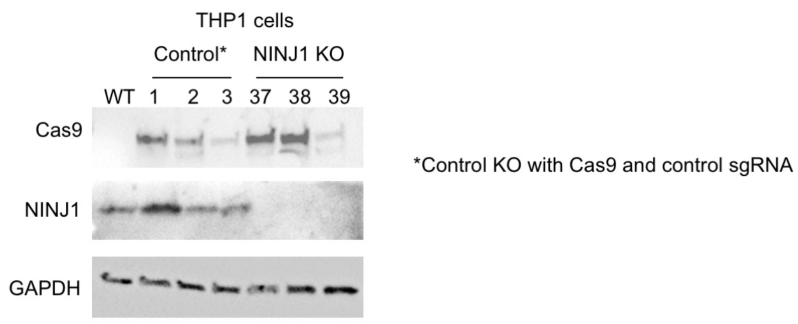


B

GenScript gRNA 6: GGCACATAGAAGGCGAAGCT

```
CTG ATG GCC AAC GCG TCC CAG CTG AAG GCC GTC GTG GAA CAG GGC CCC AGC TTC GCC TTC TAT GTG CCC CTG GTG
 L   M   A   N   A   S   Q   L   K   A   V   V   E   Q   G   P   S   F   A   F   Y   V   P   L   V
57                                                                                                   81

Clone 37:
CTG ATG GCC AAC GCG TCC CAG CTG AAG GCC GTC GTG GAA CAG GGC CCC AGC TTC GCC TTC TAT GTG CCC CTG GTG

(24 bp deletion/22 bp deletion)
CTG ATG GCC AAC GCG TCC CAG CTG AAG GCC GTC GTG GAA CAG GGC CCC AG- --- --- --- --- --- --- --- GTG
CTG ATG GCC AAC GCG TCC CAG CTG AAG GCC G-- --- --- --- --- --- --- --- -CC TTC TAT GTG CCC CTG GTG

Clone 38:
CTG ATG GCC AAC GCG TCC CAG CTG AAG GCC GTC GTG GAA CAG GGC CCC AGC TTC GCC TTC TAT GTG CCC CTG GTG

(21 bp deletion/51 bp deletion)
GCG AT- --- --- --- --- --- --- --G GCC GTC GTG GAA CAG GGC CCC AGC TTC GCC TTC TAT GTG CCC CTG GTG
CTG ATG GCC AAC GCG --- --- --- --- --- --- --- --- --- --- --- --- --- --- --- --- --- CCC CTG GTG

Clone 39:
CTG ATG GCC AAC GCG TCC CAG CTG AAG GCC GTC GTG GAA CAG GGC CCC AGC TTC GCC TTC TAT GTG CCC CTG GTG

(1 bp deletion/11 bp deletion)
CTG ATG GCC AAC GCG TCC CAG CTG AAG GCC GTC GTG GAA CAG GGC CCC AGC -TC GCC TTC TAT GTG CCC CTG GTG
CTG ATG GCC AAC GCG TCC CAG CTG AAG GCC GTC GTG GAA CAG G-- --- --- --- GCC TTC TAT GTG CCC CTG GTG
```

**Figure S4. Validation of NINJ1 knockout THP-1 cells**

(A) Western blot of anti-cas9, anti-NINJ1 and anti-GAPDH (loading control) for WT, control gRNA cells and single NINJ1 knockout clones.

(B) Genomic sequencing for NINJ1 KO clones (Clones 37, 38 and 39).

Related to Figure 4.
